# USP28 and SPINT2 mediate cell cycle arrest after whole genome doubling

**DOI:** 10.1101/2020.09.18.303834

**Authors:** Katarzyna Seget-Trzensiok, Sara Vanessa Bernhard, Christian Kuffer, Dragomir B Krastev, Mirko Theis, Kristina Keuper, Jan-Eric Boekenkamp, Maik Kschischo, Frank Buchholz, Zuzana Storchova

**Author notes:** Corresponding author: Zuzana Storchova.

## Abstract

Tetraploidy is frequent in cancer and whole genome doubling shapes the evolution of cancer genomes, thereby driving the transformation, metastasis and drug resistance. Yet, human cells usually arrest when they become tetraploid due to p53 activation that leads to CDKN1A expression, cell cycle arrest, senescence or apoptosis. To uncover the barriers that block proliferation of tetraploids, we performed an RNAi mediated genome-wide screen in a human cancer cell line. We identified 140 genes whose depletion improved survival of tetraploids and characterized in depth two of them: SPINT2 and USP28. We show that SPINT2 is a general regulator of CDKN1A, regulating its transcription via histone acetylation. By mass spectrometry and immunoprecipitation, we show that USP28 interacts with NuMA1 and affects centrosome clustering. Moreover, tetraploid cells accumulate DNA damage and loss of USP28 reduces checkpoint activation. Our results indicate three aspects that contribute to survival of tetraploid cells: i) increased mitogenic signaling and reduced expression of cell cycle inhibitors, ii) the ability to establish functional bipolar spindle, and iii) reduced DNA damage signaling.

## Introduction

Recent progress in cancer genome analysis has exposed the complexity of genomic alterations during tumorigenesis. One of the frequent types of cancer-associated genomic alterations is whole-genome doubling (WGD) that arises due to defects in mitosis and cytokinesis as well as via cell-cell fusion or chromosome endoreduplication (Davoli et al., 2011, Lens et al., 2019, Storchova et al., 2004). Computational analysis of human exome sequences from ∼4,000 human cancers suggest that approximately 37% of all tumors have undergone at least one whole-genome-doubling event at some point during tumorigenesis (Bielski et al., 2018, Dewhurst et al., 2014, Zack et al., 2013); the frequency of WGD rises to 56 % in metastasis (Priestley et al., 2019). A passage through tetraploid intermediate facilitates chromosomal instability and contributes to increased tumorigeneity, metastasis formation, drug resistance and is associated with a reduced chance of disease-free survival (Bielski et al., 2018, Dewhurst et al., 2014, Galofre et al., 2020, Kuznetsova et al., 2015, Wangsa et al., 2018). Tetraploid cells also show an increased tolerance to further chromosomal abnormalities (Dewhurst et al., 2014, Kuznetsova et al., 2015). However, experimental induction of tetraploidy is not well tolerated in mammalian cells, which poses a question as of how can cells survive and tolerate tetraploidy (Ganem et al., 2007, Lens et al., 2019). Identification of pathways that limit the survival of tetraploid cells may therefore improve our understanding of cancerous processes and pave the way for novel cancer treatments.

*In vitro*, the proliferation of tetraploid cells arising from whole genome doubling is limited at two control points (Fig 1A). First, induced cytokinesis failure may trigger a cell cycle arrest immediately in the following G1 phase by stabilizing the tumor suppressor protein p53 and elevating the expression of its downstream target CDKN1A (Lanni et al., 1998), (Andreassen et al., 2001). This is often observed in non-transformed cells, such as mouse embryonic fibroblasts (MEFs) or in the human immortalized RPE1 cell line. A genome-wide screen in RPE1 revealed that the cytoskeletal stress caused by extra centrosomes in binucleated tetraploids activates the Hippo tumor suppressor pathway (Ganem et al., 2014). The Hippo pathway kinases LATS1 and LATS2 then inhibit the MDM2 E3 ligase, thereby stabilizing p53 (Aylon et al., 2006, Ganem et al., 2014). The G1 arrest immediately after cytokinesis failure can be also bypassed by enhanced cytokine signaling (Ganem et al., 2014, Vittoria et al., 2018).

**Figure 1.**
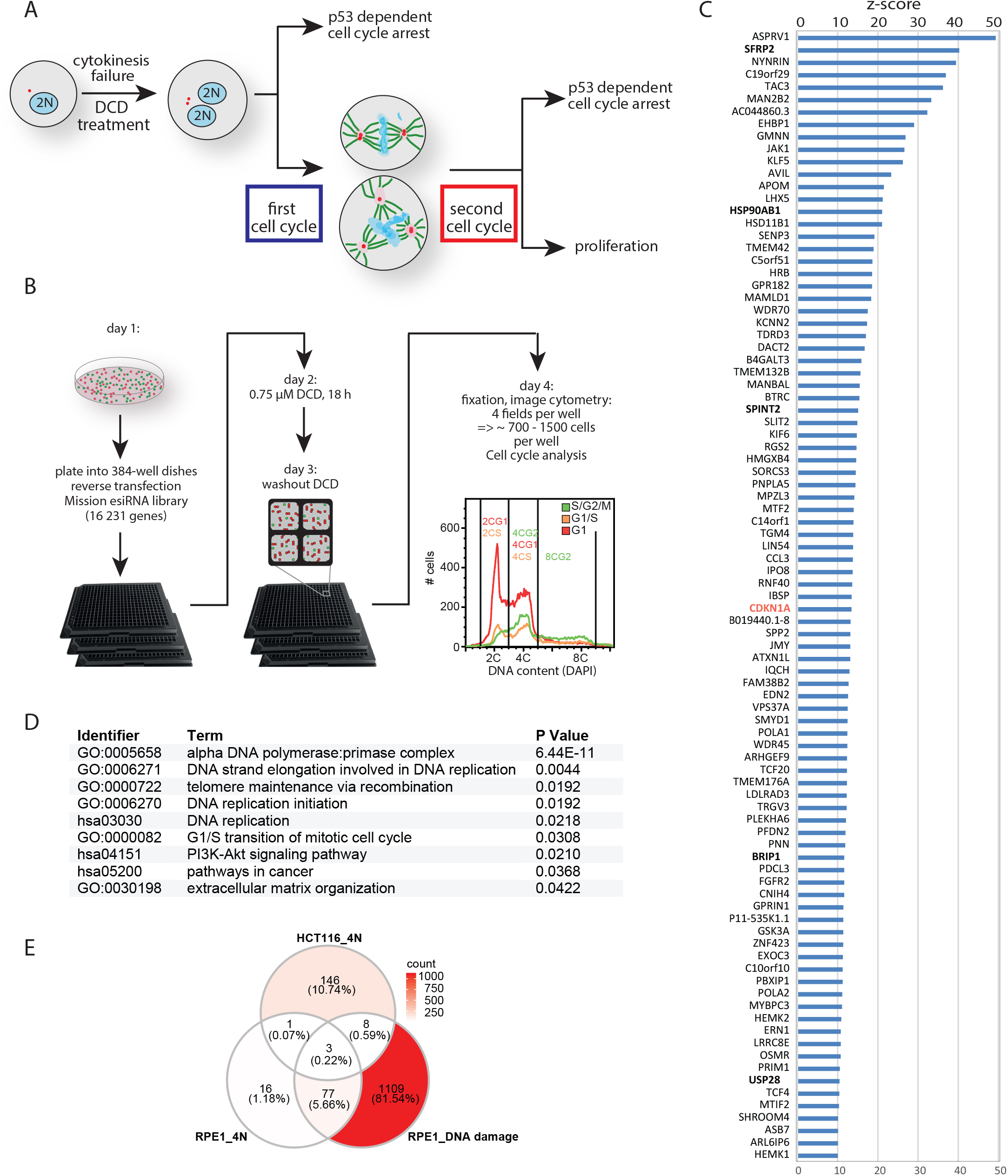
Genome-wide screen identifies novel factors involved in proliferation of cancerous tetraploid cells. A. Schematic depiction of the possible fates of human cells upon cytokinesis failure. B. Schematic depiction of the microscopy-based esiRNA screen set up. C. Top candidates identified in the screen (Z-score>10, high significance). D. Pathway enrichment analysis identifies nine significantly enriched pathways among the screen hits. E. Venn diagram of overlap between hits in gene depletion screens for tetraploid HCT116 survival improvement (HCT116_4N) and for RPE1 cell cycle arrest escape after induced tetraploidization (RPE1_4N) and DNA damage (RPE_DNA damage) (Ganem et al., 2014).

However, mammalian cells can often escape the G1 arrest immediately after tetraploidization and enter the cell cycle (Dewhurst et al., 2014, De Santis Puzzonia et al., 2016, Kuffer et al., 2013, Lv et al., 2012, Wangsa et al., 2018, Wong et al., 2005). The subsequent mitoses are often aberrant due to the presence of extra centrosomes and doubled genetic material (Fig 1A). Multipolar mitosis or erroneous bipolar mitosis may also trigger a p53-dependent arrest in the subsequent G1 phase (Ganem et al., 2009, Kuffer et al., 2013), but little is known about the involved molecular mechanisms. Abnormal tetraploid mitosis increases oxidative damage and knock down of ATM improves subsequent proliferation (Kuffer et al., 2013). Another report demonstrates that an increase in the number of mature centrosomes activates the PIDDosome complex. This in turn leads to Caspase-2-mediated MDM2 cleavage, p53 stabilization, and p21-dependent cell cycle arrest (Fava et al., 2017). Tetraploidy may also induce replication stress and activate the ATR-Chk1-signalling (De Santis Puzzonia et al., 2016, Pedersen et al., 2016, Wangsa et al., 2018). Increased expression of cyclin D1 and D2 was reported in surviving tetraploid cells, suggesting that the G1 arrest can be bypassed by enhancing the expression of G1 cyclin A (Crockford et al., 2017, Potapova et al., 2016). Taken together, the findings imply that there are several routes to arrest after aberrant tetraploid mitosis and that we are far from understanding the molecular mechanisms that allow to escape this fate.

The meta-analysis of cancer genomes suggests that whole-genome doubling occurs not only in early precancerous lesions, but frequently also after transforming mutations of cancer driver genes, for example as an event facilitating metastases (Bielski et al., 2018, Priestley et al., 2019). The factors that limit proliferation of transformed tetraploid cells may differ from factors important for arrest of non-transformed ones. Therefore, we performed an RNAi based high throughput screen for factors limiting proliferation after cytokinesis failure in the near-diploid, but transformed cell line HCT116. These cells do not arrest in the first G1 after becoming tetraploid, but enter the next cell cycle and arrest later, after subsequent aberrant tetraploid mitoses (Fig 1A, (Kuffer et al., 2013). Using siRNA and CRISPR/Cas9 techniques, we evaluated in detail the effect of two of the identified candidates, SPINT2 and USP28. We demonstrate that SPINT2 affects mitogenic signaling as well as CDKN1A expression, and its loss enables to bypass the G1-arrest upon various cellular stresses. Additionally, we show that proliferating tetraploid cells quickly accumulate DNA damage, and loss of USP28 reduces the checkpoint response, thus allowing proliferation of tetraploid cells. Our findings demonstrate that the context of tetraploidization is important for cellular response and survival, and suggest new mechanisms that enable tetraploid proliferation.

## Results

### High throughput genetic screen to identify factors involved in arresting the tetraploid cells

Proliferation of tetraploid cells is inhibited either immediately after cytokinesis failure or after the first tetraploid mitosis that is usually erroneous due to multiple centrosomes (Fig 1A). To determine factors that control the proliferation of transformed tetraploid cells, we used HCT116, a pseudo-diploid (45,X) transformed human p53-positive colorectal cancer cell line. The cells were additionally modified with Fluorescent Ubiquitination-based Cell Cycle Indicator (FUCCI, (Sakaue-Sawano et al., 2008)). The FUCCI G_1_ sensor consists of the N-terminus of Cdt1 fused to mKO2 (Kyoto Orange fluorescent protein) that is expressed in G1 and degraded in S, G_2_ and M phase by the SCF complex, and the FUCCI G_2_ sensor based on N-terminus of Geminin fused to mAG (Azami Green fluorescent protein) that is degraded by the APC/C complex at the end of mitosis. To generate tetraploid cells, HCT116 cells were treated 18 h with dihydrocytochalasin D (DCD), an inhibitor of actin polymerization that prevents cytokinesis. This treatment results in a mixed population of approximately 60-70 % binucleated tetraploid and 30-40 % diploid HCT116 (Kuffer et al., 2013). Under these conditions, more than 80 % of both binucleated tetraploids and diploids entered the next S phase and progressed to mitosis. Whereas diploid cells proliferate normally, the subsequent mitoses of tetraploid cells are often erroneous, and about 50 % of HCT116 cells arrest shortly after the second tetraploid mitosis (Kuffer et al., 2013). To identify factors that influence this process, we performed a genome-scale RNAi screen in HCT116 where the cells were subjected to DCD treatment. In our settings, the diploid and tetraploid populations can be distinguished by DNA content in combination with the FUCCI sensor by image analysis of the mixed population (Fig 1B). This strategy provided two main advantages. First, by avoiding FACS sorting of tetraploid and diploid population, we minimized the manipulations with the cells and possible artifacts. Second, by comparing the effects of knockdown in diploids and tetraploids side-by-side in the same experiment, we directly identified the tetraploidy-specific factors.

To identify factors that control proliferation of tetraploids, we used an esiRNA library targeting 16,231 genes in the human genome (Kittler et al., 2007, Krastev et al., 2011, for full details about the screening strategy see Supplementary Information). The primary screen identified 432 genes whose depletion improved proliferation of tetraploid cells; we name this category a TP53-like class, since knock down of these genes improves proliferation of tetraploid cells similarly to knock down of TP53. Additionally, we identified 1150 genes whose depletion killed tetraploid cells (CK, personal communication). Next, we performed a validating screen of primary TP53-like hits and confirmed 157 genes from the primary screen (Supplementary Table 1). We then calculated the Z-scores and selected genes with a Z-score >10, obtaining a high confidence group of 90 genes (Fig 1D). Among the strongest hits was CDKN1A (p21), a downstream target of p53 that promotes cell cycle arrest. CDKN1A was previously found to inhibit tetraploid proliferation, thus validating the overall strategy (Vittoria et al., 2018). Pathway enrichment analysis of the hits by DAVID (Dennis et al., 2003) determined statistically significant enrichment of the pathways related to DNA replication (e.g. “DNA polymerase:primase complex”, “DNA replication initiation”), “G1-S transition of mitotic cell cycle”, “PI3k-Akt signaling pathway”, “Pathways in cancer” and “Extracellular matrix organization” (Fig 1D, Supplementary Table 2). Comparing the list of genes identified in our screen with previous results revealed only minor overlap with genes that were found to limit proliferation of tetraploid RPE1 (Ganem et al., 2014). In fact, we found a larger overlap with factors required for survival after DNA damage (Fig 1E, Supplementary Table 3). For subsequent analysis, we selected six candidates that represent each category and were also previously linked to colorectal cancer.

### SPINT2 and USP28 regulate proliferation of tetraploid cells

For validation and functional follow up, we selected SFRP2 (Secreted Frizzled Related Protein 2), coding for a soluble modulator of WNT signaling, HSP90AB1 (Heat Shock Protein 90 Alpha Family Class B Member 1) coding for molecular chaperone, CCDC6 (Coiled-Coil Domain Containing 6), and uncharacterized putative tumor suppressor, SPINT2 (Serine Peptidase Inhibitor, Kunitz Type) coding for hepatocyte growth factor inhibitor, BRIP1 (BRCA1 Interacting Protein C-Terminal Helicase 1), a breast cancer associated gene coding for a protein involved in DNA repair and USP28 (Ubiquitin Specific Peptidase 28), a deubiquitinase involved in DNA damage response checkpoint and MYC proto-oncogene stability (Fig 1D). To validate the selected candidates, we treated the cells with siRNA of respective proteins, followed by DCD to induce cytokinesis failure. Samples were collected 24, 30 and 48 h after the DCD washout. 30 min before collecting the samples, the cells were treated for 30 min with EdU (5-ethynyl-2’-deoxyuridine) that was incorporated into the replicating DNA to determine the fraction of proliferating cells (Fig 2A, EV1A). Subsequent flow cytometry quantified the proportion of proliferating diploids and tetraploids and their cell cycle phase (Fig 2B, EV1A). Treatment with respective siRNA depleted on average 80 % of the candidate proteins (Fig EV1B). Knock down of SPINT2 and USP28 improved tetraploid proliferation, thus confirming their role in cellular response to tetraploidy. Depletion of the other factors did not affect proliferation of tetraploids and they were excluded from further analysis (Fig 2C). No effect was observed in diploids (Fig EV1C). Analysis in RPE1 cells confirmed the role of p53 and SPINT2 in tetraploid proliferation that have been shown previously, but we observed no effect of USP28 on survival of tetraploid RPE1 cells (Fig 2D). Time-lapse live cell imaging further validated that depletion of SPINT2 and USP28 increased the number of tetraploid HCT116 that entered second mitosis after cytokinesis failure (Fig 2E). Finally, we used CRISPR/Cas9 to knock out SPINT2 and USP28 in HCT116 cells (Fig EV1D, E). The knock out cell lines showed improved proliferation upon induced cytokinesis failure that was abolished upon transfection with plasmid carrying the wild type gene (Fig 2F-H, EV1F, G). We conclude that SPINT2 and USP28 negatively affect proliferation of newly arising tetraploid cells.

**Figure 2.**
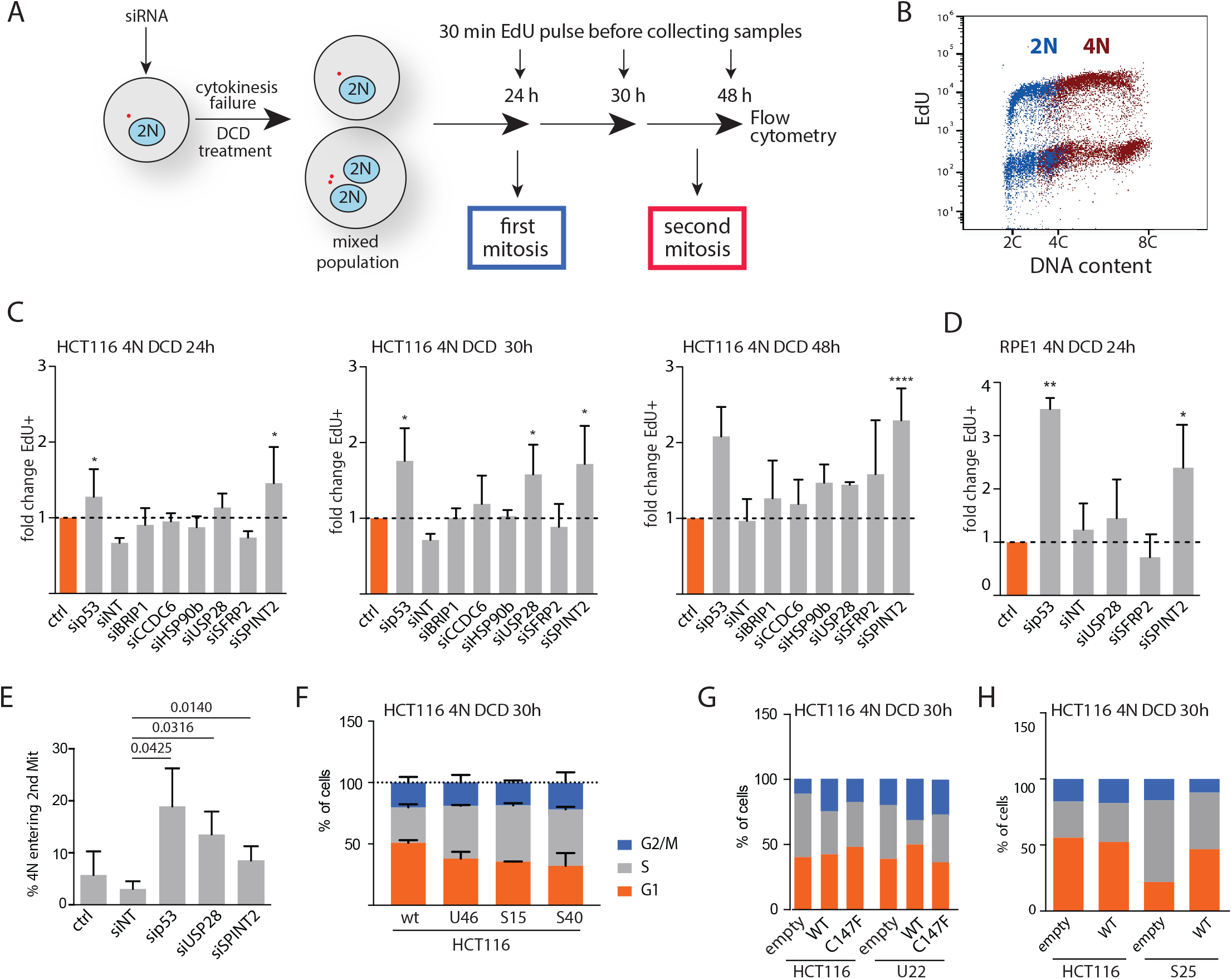
Validation of the identified targets confirms the role of USP28 and SPINT2 in proliferation of transformed tetraploid cells. A. Schematics of the experimental set up to validate the targets. B. Cell cycle profile obtained by flow cytometry of EdU labeled cells after cytokinesis failure. The EdU labeling allows to identify proliferating cells; the staining with PI allows to distinguish the cells according to their DNA content. Cell categories: diploid in G1: 2C_G1, diploid in S, G2, M: 2C_S, tetraploid in G1: 4C_G1, tetraploid in S: 4C_S and tetraploid in G2 and M: 8C_G2). C. Fold change of proliferating cells 24 h, 30 h and 48 h after cytokinesis failure in HCT116 treated with siRNA against respective genes or with a non-targeting control (siNT) compared to an untreated control (ctrl). siRNA against p53 was used as a positive control. D. Fold change of proliferating cells 24 h after cytokinesis failure in RPE1 treated with siRNA against respective genes or with a non-targeting control compared to an untreated control. siRNA against p53 was used as a positive control. E. Percentage of cells entering 2^nd^ tetraploid mitosis after cytokinesis failure in HCT116 treated with siRNA against respective genes or with a non-targeting control. siRNA against p53 was used as a positive control. Time-laps live cell imaging, 8 min time frame, 72 h, 20x air objective. At least three independent experiments, each at least 40 cells per condition, were analyzed. F. Cell cycle distribution analysis in tetraploids generated from *USP28* and *SPINT2* knockout clones (U46 and S15, S40, respectively) using flow cytometry. Knockouts were generated with CRISPR/Cas9 in HCT116. Samples were analyzed 30 h after a release from the DCD treatment. G. Cell cycle distribution analysis in tetraploids generated from *USP28* knockout clone U22 after transfection with a control empty plasmid, plasmid with wildtype *USP28* and a plasmid containing a mutant *USP28* lacking the deubiquitinase active site (C147F)). Representative plot of three biological experiments is shown. H. Cell cycle distribution analysis in tetraploids generated from *SPINT2* knockout clone S25 after transfection with a control empty plasmid or plasmid with wildtype *SPINT2*. Representative plot of three biological experiments is shown.

### SPINT2 and USP28 act independently of Hippo signaling or PIDDosome activation

Formation of tetraploid cells activates the p53 response and induces the expression of p21 in most human cell lines (Andreassen et al., 2001). We therefore tested whether the depletion of SPINT2 and USP28 diminishes the p53 activity, thereby facilitating the growth of tetraploid cells. While the p53 levels were not affected by SPINT2 and USP28 depletion, the expression of p21 was strongly reduced upon SPINT2 depletion and partly also upon USP28 depletion (Fig 3 A-C). Activation of Hippo signaling and PIDDosome were previously shown to control the survival of tetraploid cells (Fava et al., 2017, Ganem et al., 2014). Since we did not identify any members of these two pathways, we asked whether they were activated in response to tetraploidy in the HCT116 cell line. Indeed, we found that MDM2 and caspase 2 became cleaved in response to cytokinesis failure in HCT116, demonstrating that PIDDosome activity increased upon whole genome doubling as previously described (Fig EV2A, B). Importantly, a depletion of SPINT2 or USP28 did not affect the MDM2 and caspase 2 cleavage (Fig EV2B). In contrast, the YAP localization was not altered and the phosphorylation of LATS2 was not increased in tetraploid HCT116, suggesting that cytokinesis failure in HCT116 cells does not activate the Hippo pathway (Fig EV2C-E). Taken together, SPINT2 and USP28 affect proliferation of tetraploid cells independently of Hippo or PIDDosome signaling.

**Figure 3.**
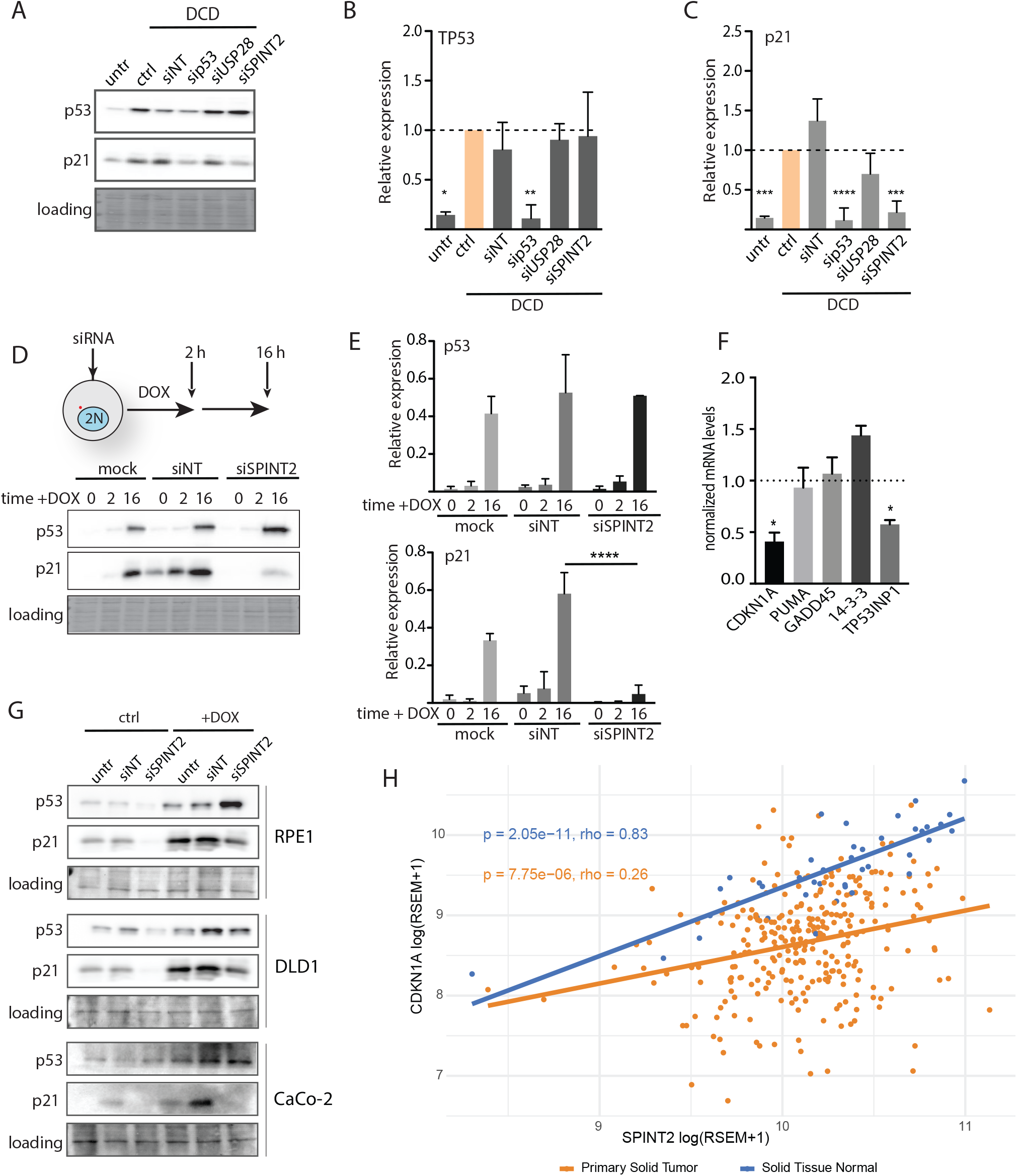
SPINT2 modulates expression of p21 independently of p53 levels. A. Expression levels of p53 and p21 upon depletion of the candidate factors in DCD treated HCT116. Ponceau staining was used as a loading control. B. C. Quantification of p53 and p21 levels upon depletion of the candidate factors in DCD treated cells. Mean and SEM of three experiments is shown, t test was used for statistical evaluation. D. Schematic presentation of the experiment and the expression of p53 and p21 in doxorubicin (DOX) treated cells depleted for SPINT2. Ponceau staining was used as a loading control. E. Quantification of p53 and p21 levels upon SPINT2 depletion in DOX treated cells. Mean and SEM are shown, t test was used for statistical evaluation. F. Normalized mRNA levels of p53 downstream targets upon SPINT2 depletion in DOX treated cells. Mean and SEM of at least three independent experiments are shown, normalized to control housekeeping gene *RPL30*. G. Immunoblots of p53 and p21 in RPE1, DLD1 and CaCO upon SPINT2 depletion. Ponceau staining was used as a loading control. H. Correlation of CDKN1A and SPINT2 mRNA expression levels in colorectal adenocarcinoma and in normal tissues. RSEM (RNA-Seq by Expectation Maximization) – quantification of the RNA abundance from RNA-seq data.

### SPINT2 regulates the expression of a subset of TP53 targets

SPINT2, a putative tumor suppressor, encodes a transmembrane protein with two extracellular protease inhibitor domains (Kunitz domains) that act on a variety of serine proteases (Friis et al., 2014). It inhibits the binding of signaling molecules, such as HGF (hepatocellular growth factor) to the c-Met receptor and thereby affects ERK, AKT and STAT3 pathways (Fig EV3A). Thus, loss of SPINT2 is predicted to enhance growth factor signaling and indeed, this was sufficient to overcome the G1 arrest in tetraploid non-cancerous RPE1 cells (Ganem et al., 2014). In diploid HCT116, the loss of SPINT2 also enhanced the activation of AKT pathway in response to growth factors (Fig EV3B). We observed that upon release of starving cells to serum-proficient medium, the phosphorylation of AKT was significantly increased and the expression of p21 nearly abolished in cells with reduced SPINT2 (Fig EV3B, C). Moreover, the nuclear levels c-Myc, a pro-proliferative transcription factor and cMET target (via STAT3) (Zhang et al., 2002) increased significantly upon SPINT2 knock down (Fig EV3D). Thus, loss of the cMET inhibitor SPINT2 bypasses the cell cycle arrest by activation of pro-proliferative factors.

A striking phenotype of SPINT2 was the reduction of CDKN1A (p21) expression despite p53 activation (Fig 3A-C), which is likely the reason for improved tetraploid proliferation. We asked whether SPINT2 depletion reduces CDKN1A expression also in response to other cellular insults that activate the p53 pathway, such as DNA damage. To this end, we treated diploid cells with the topoisomerase II inhibitor doxorubicin (DOX). This causes DNA damage that activates the p53-mediated response and results in a stabilization of p53 and increased p21 levels. Strikingly, the depletion of SPINT2 abolished CDKN1A activation upon DOX treatment, while the p53 levels were not affected, similarly as in the response to tetraploidy (Fig 3D-F).

The transcription factor p53 activates the expression of multiple different targets in response to cellular stress. Using rt-PCR, we evaluated the effect of SPINT2 depletion on expression of p53 targets upon treatment with DOX. The expression of factors required for apoptosis, such as GADD45 or PUMA, were not affected by SPINT2 depletion. In contrast, expression of factors that promote cell cycle arrest, CDKN1A and TP53INP1, were reduced in response to DNA damage when SPINT2 was depleted (Fig 3F). The reduced CDKN1A expression upon SPINT2 depletion was observed also in RPE1, DLD1 and CaCo2 cell lines (Fig 3G). Finally, data analysis of gene expression from The Cancer Genome Atlas database (TCGA) revealed a strong correlation between the CDKNA1A and SPINT2 expression levels in healthy samples, and a weak correlation in cancers samples, where SPINT2 is frequently mutated (Fig 3H) (Dong et al., 2010). Thus, SPINT2 is a general regulator of the CDKN1A expression.

To investigate how SPINT2 affects CDKN1A expression, we performed chromatin immunoprecipitation of p53 on the defined regulatory elements of the CDKN1A promoter. Strikingly, the binding of TP53 to the CDKN1A promoter is lost upon SPINT2 depletion in DOX or DCD treated cells (Fig 4A). This is not due to the lack of p53 in the nucleus (Fig 4B). Histone acetylation within the CDKN1A promoter region is essential for its expression (Lagger et al., 2003). We tested whether inhibition of histone deacetylases (HDAC) restores the expression of the CDKN1A in cells lacking SPINT2. To this end, we used the knock out HCT116 cells lacking SPINT2. We treated the cells with trichostatin A (TSA), a potent HDAC inhibitors, simultaneously with doxorubicin. Strikingly, the loss of SPINT2 no longer suppressed the expression of p21 when histone deacetylases were inhibited (Fig 4C-E). We conclude that SPINT2 is a previously unidentified regulator of CDKN1A expression in human cells that may modulate the histone modification within the CDKN1A promoter.

**Figure 4.**
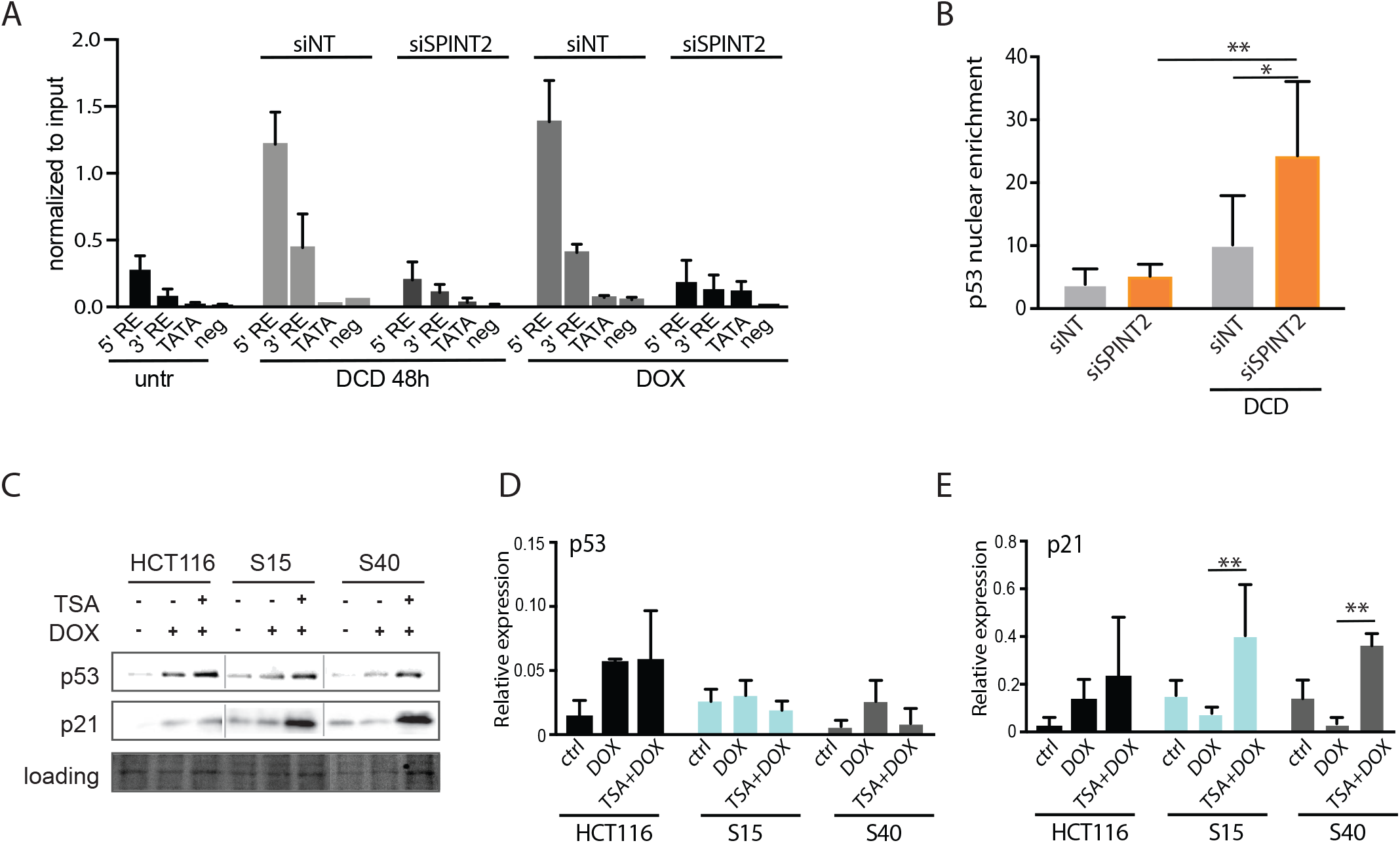
The effect of SPINT2 on CDKN1A (p21) expression is mediated via promotor acetylation. A) Chromatin immunoprecipitation of p53 with the defined regulatory elements within the *CDKN1A* promoter. B) Nuclear enrichment of p53 after siRNA knockdown of SPINT2. NT: non-targeting siRNA control. Mean and SD of three independent experiments is shown, t test, * p< 0.05; ** p < 0.01. C) Representative immunoblot of p53 and p21 in HCT116 and two SPINT2 knockout clones S15 and S40 after TSA and doxorubicin (DOX) treatment. Ponceau staining was used as a loading control. D) E) Quantification of p53 and p21 levels from three independent experiments normalized to the loading control. Mean with SD is shown, t test. * p< 0.05; ** p < 0.01.

### Loss of USP28 affects centrosome clustering

USP28 was recently identified as a factor that together with 53BP1 and TRIM37 stabilizes p53 in response to centrosomal stress or after extended duration of mitosis in RPE1 cells (Fong et al., 2016, Lambrus et al., 2016, Meitinger et al., 2016). No effect of USP28 was observed after cytokinesis failure in RPE1. This is in line with our observations (Fig 2D). The function of USP28 after tetraploidization in HCT116 differs from its effect after centrosomal depletion, since 53BP1 did not influence proliferation of tetraploid HCT116 (Fig EV4A). Moreover, although the mitosis in tetraploids generally takes longer than in diploids, it only rarely exceeds 90 min (Fig EV4B). Because most RPE1 cells arrests in the G1 immediately following the cytokinesis failure, while HCT116 progresses through 2-3 cell cycles before arresting, we hypothesized that USP28 is required for the cellular response of proliferating HCT116 tetraploids, which cannot be observed in the primary cell lines.

To elucidate the function of USP28 in response to tetraploidy, we performed immunoprecipitation (IP) of proteins interacting with USP28 followed by mass spectrometry. Specifically, we analyzed HCT116 with and without DCD treatment, looking for interactors 24 h after cytokinesis failure. Strikingly, we found several proteins whose presence upon co-IP with USP28 was significantly increased in DCD-treated samples (Fig 5A). Among them were mediator of DNA damage checkpoint 1 (MDC1) that has been already previously identified as a USP28 interactor upon DNA damage (Zhang et al., 2006). This interaction is required for full activation of DNA damage checkpoint. Interestingly, we also found an interaction with Nuclear mitotic apparatus protein 1 (NuMA1) that is involved in spindle apparatus and microtubule functions and has not been previously related to USP28. The co-IP of USP28 and NuMA1 was specific for tetraploid cells, as confirmed by a pull down followed by immunoblotting (Fig 5B). Strikingly, both USP28 and NuMA1 also colocalized on centrosomes (Fig 5C, Fig EV4 C).

**Figure 5.**
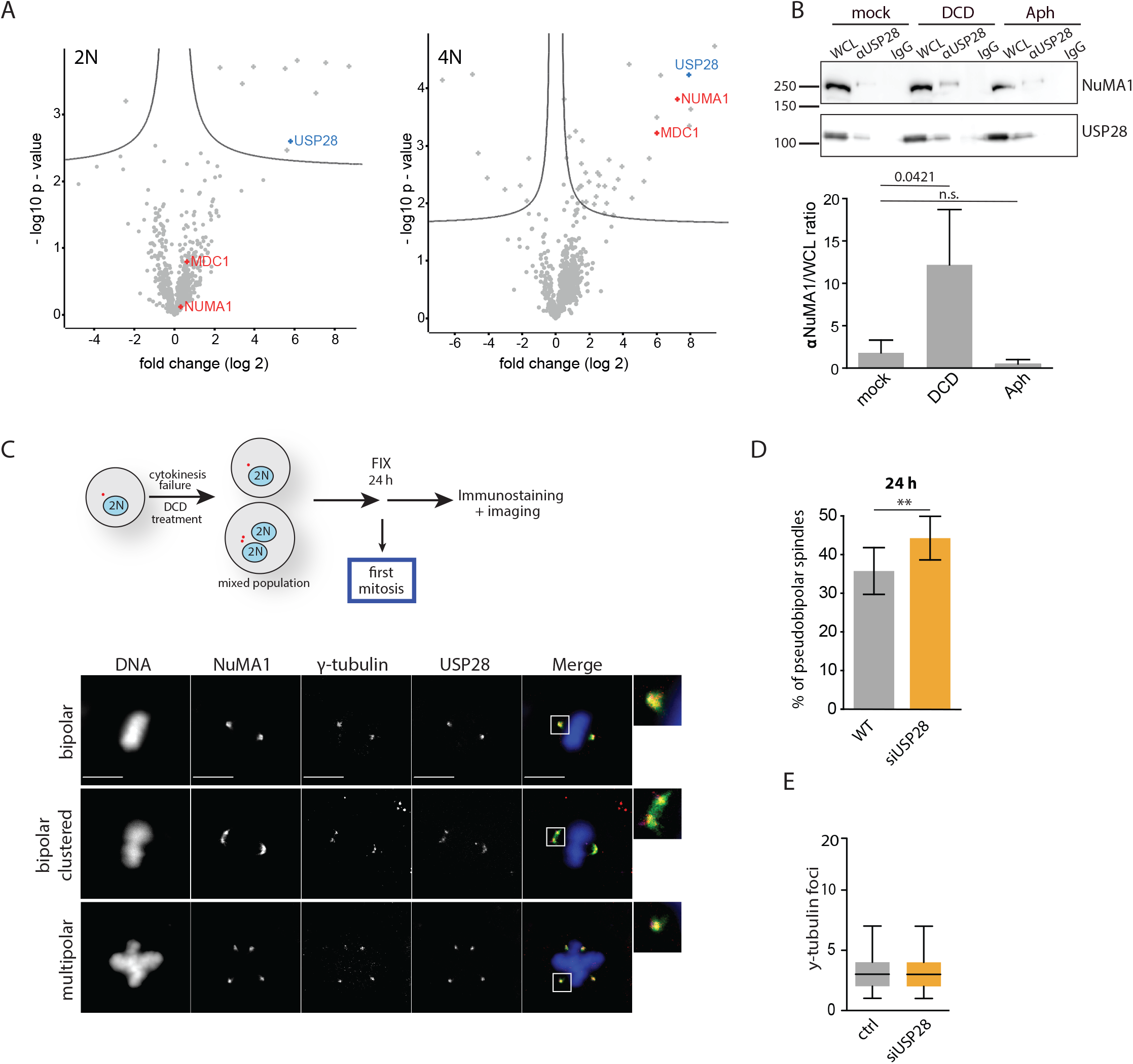
USP28 affects centrosome clustering in proliferating tetraploids. A) Volcano blot showing the interaction partners of USP28 (blue) in diploid and tetraploid HCT116. The interactors NuMA1 and MDC1 are marked in green. Complete results are in Supplementary Table 4. B) Immunoblot of USP28 and NuMA1 after co-immunoprecipitation of USP28 using magnetic beads (*upper panel*). Ratio between NuMA1 and respective whole-cell lysates (*lower panel*). Samples were treated with DCD and aphidicolin (Aph), respectively. C) Schematics of the experiment and immunofluorescence pictures in HCT116 γ-tubulin mRuby stained for USP28 and NuMA1 showing localization of USP28 and NuMA1 in relation to γ-tubulin. DNA was stained using DAPI. Scale bars: 10 μm. D) Percentage of pseudo-bipolar spindles in tetraploid HCT116 γ-tubulin mRuby and after USP28 siRNA knockdown. Cells were fixed 24h after DCD washout and analyzed via immunofluorescence pictures. T test was used for statistical evaluation. E) Analysis of centrosome numbers in tetraploid HCT116 γ-tubulin mRuby and after USP28 siRNA knockdown. T test was used for statistical evaluation.

One of the well-recognized functions of NuMA1 is its involvement in clustering of supernumerary centrosomes (Quintyne et al., 2005). We asked whether the USP28-NuMA1 interaction might regulate the centrosome clustering in tetraploid cells. Mitotic figures in the first tetraploid mitosis with and without USP28 revealed that HCT116 tetraploid cells clustered the spindle poles more efficiently in the absence of USP28 than control cells (in average 45 %, compared to 35 %, Fig 5D), while the numbers of centrosomes were not altered (Fig 5E). Since pseudobipolar mitosis leads to less cellular death than multipolar mitosis (Ganem et al., 2009), this increased centrosomal clustering might improve viability of tetraploid cells. USP28 is a deubiquitinase and regulates stability of several proteins. However, we did not observe any difference in NuMA1 abundance upon USP28 depletion and thus the mechanism of the effect of USP28 on centrosome clustering remains unclear (Fig EV4 D, E) Taken together, USP28 affects clustering of centrosomes in tetraploid cells in cooperation with NuMA1, thereby decreasing the probability of detrimental multipolar mitoses.

### Increased DNA damage in tetraploid cells

Another interesting interactor of USP28 specifically enriched in tetraploid cells was MDC1, a facilitator of DNA damage response and checkpoint activation (Zhang et al., 2006). Previously, it has been suggested that WGD triggers DNA damage in proliferating cells, as has been shown in Drosophila or U2OS (Nano et al., 2019, Pedersen et al., 2016). Thus, depletion of USP28 may contribute to tetraploid proliferation by reducing the checkpoint activation upon DNA damage that progressively accumulates in proliferating tetraploid cells. Evaluation of DNA damage and replication stress markers revealed an increased phosphorylation of RPA32 at Ser33 and, with a delay, at Ser4/8 (targets of ATR and DNA-PK, respectively (Marechal et al., 2015), Fig 6A). The yH2AX signal that marks DNA damage also increased during the 48 h time course after cytokinesis failure, as well as the phosphorylation of CHK1 and accumulation of p53 and p21 (Fig 6A, B). Strikingly, depletion or knock out of USP28 diminished the checkpoint activation (Fig 6A, B). The time course demonstrated that the DNA damage accumulates only when the tetraploid cells enter cell cycle, after at least one round of replication. Thus, loss of USP28 alleviates the checkpoint activation upon DNA damage, thereby enhancing the proliferation of tetraploid cells.

**Figure 6.**
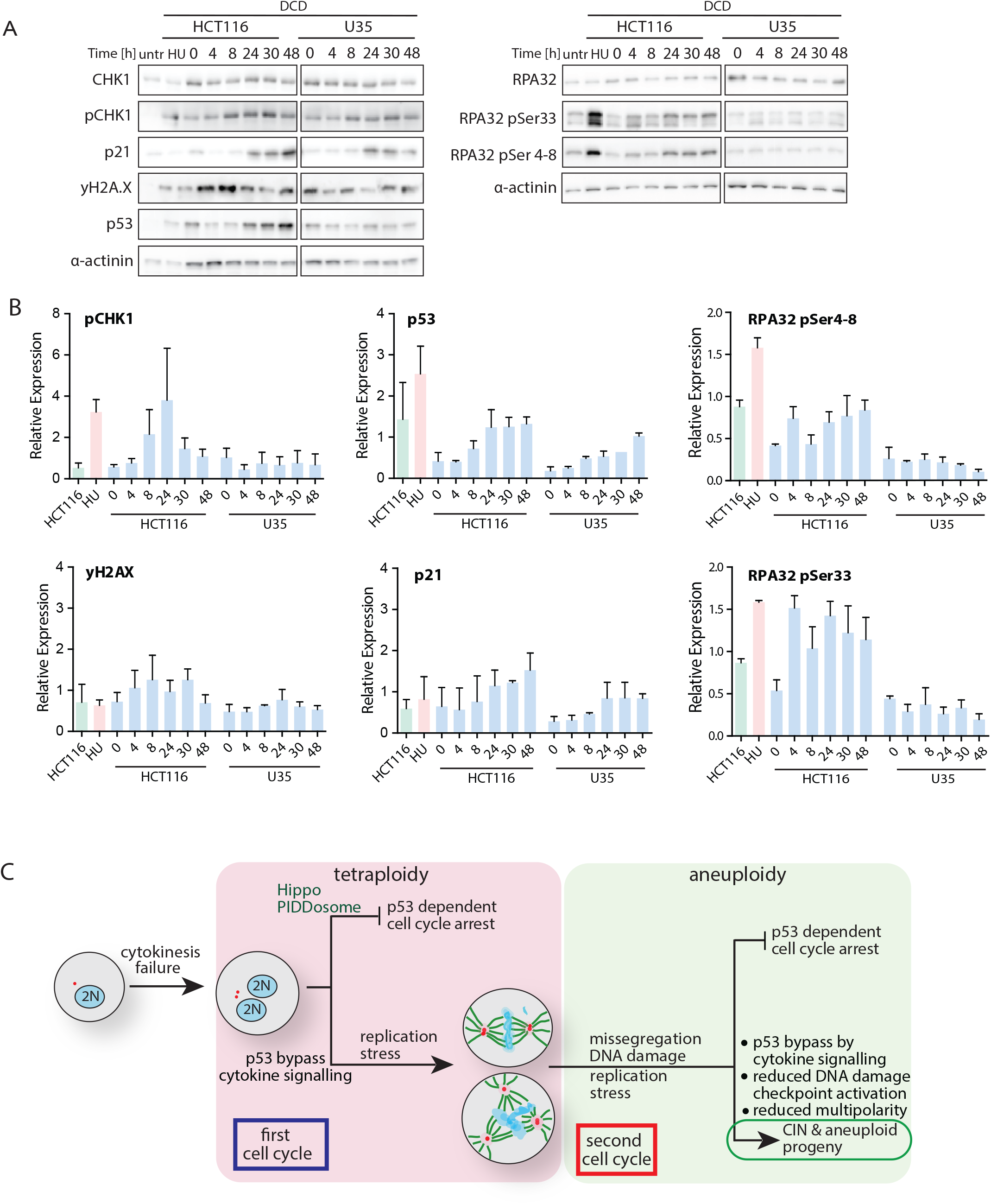
The DNA damage signaling in tetraploid cells is reduced upon loss of USP28. A) Immunoblot showing the time course of different DNA damage associated proteins after DCD treatment in HCT116 (left) and after USP28 depletion (right). Ponceau staining was used as a loading control. B) Immunoblot of different DNA damage associated proteins in tetraploid HCT116 and after USP28 knockout. α-actinin was used as a loading control. C) Relative expression of DNA damage associated proteins in DCD treated HCT116 with and without USP28. Values were normalized to the loading control. D) Schematic depiction of the cellular fate after cytokinesis failure.

## Discussion

Whole genome doubling shapes the evolution of cancer and drives transformation, metastasis and drug resistance. Increased ploidy has been documented in nearly 37 % of cancers and is even more prevalent in advanced metastases, where it can be found in 56 % of the cases (Bielski et al., 2018, Priestley et al., 2019). But human non-transformed cells usually arrest after becoming tetraploid, which raises the question how proliferating tetraploid populations arise. Recent genomic analysis indicates that tetraploidization usually occurs in pre-cancerous lesions following a p53 inactivation or other permissive mutation (Bielski et al., 2018). We performed a genome-scale screen for factors enabling proliferation of transformed cells, in which we induced cytokinesis failure. These cells do not arrest immediately after becoming tetraploid, but rather progress through the next cell cycles and subsequently become trapped in an irreversible arrest due to accumulation of genomic abnormalities. We show that three factors become essential for survival of tetraploid cells in this context: increased mitogenic signaling and reduced expression of cell cycle inhibitors, the ability to establish bipolar spindle and reduced DNA damage signaling (Fig 6C).

Using an esiRNA library, we identified 140 genes whose depletion improved proliferation of tetraploid cells (Fig 1). Validation of selected candidates confirmed USP28 and SPINT2 as factors that negatively regulate proliferation of tetraploids (Fig 2A-D). Both factors act independently of the HIPPO and PIDDosome pathways (Fig EV2) that were previously identified to inhibit tetraploid proliferation (Fava et al., 2017, Ganem et al., 2014). Our results demonstrate that the cellular response to tetraploidy is cell type and context dependent and a complete picture of molecular processes affected by tetraploidy is still missing.

SPINT2, a putative tumor suppressor, was identified as a factor limiting proliferation of tetraploids not only in HCT116 cells, but also in RPE1 cell line (Ganem et al., 2014). SPINT2 encodes an inhibitor of growth factor signaling and, thus, the increased proliferation rates upon SPINT2 loss might be explained by overactivation of growth factor signaling, which in turn leads to a bypass of a p53-dependent cell cycle arrest (Friis et al., 2014, Ganem et al., 2014). In tetraploid cells, both activation of mitogenic signaling by loss of SPINT2 and overexpression of c-myc and cyclin D1 or D2 (downstream targets of the mitogenic signaling) lead to increased proliferation (Crockford et al., 2017, Ganem et al., 2014, Potapova et al., 2016). Thus, the cell cycle arrest of tetraploid cells might be bypassed by regulation of cell cycle inhibitors and activators. We found that SPINT2 acts as a general modulator of CDKN1A transcription (Fig 3, 4). Depletion of SPINT2 reduced CDKN1A transcription, via epigenetics regulation, upon induced cytokinesis failure as well as upon induction of DNA damage. In agreement with this observation we found that the expression levels of the SPINT2 and CDKN1A correlate in human cells. Since SPINT2 is a putative tumor suppressor in some cancer types, our findings suggest the mechanism of its tumor suppressing activity.

Loss of USP28, a deubiquitinase, also improves proliferation of tetraploid HCT116. USP28 was recently identified as a key factor affecting the proliferation upon centrosome loss in RPE1 cells (Fong et al., 2016; Lambrus et al., 2016; Meitinger et al., 2016), but not after cytokinesis failure (Meitinger et al., 2016). USP28 also interacts with 53BP1 to arrest cells upon centrosome loss and prolonged mitosis (Fong et al., 2016; Knobel et al., 2014; Lambrus et al., 2016; Meitinger et al., 2016), however, 53BP1 plays no role in proliferation of tetraploid HCT116. We propose that the observed difference between RPE1 and HCT116 is due to a different timing of the cell cycle arrest after tetraploidization. While primary cells mostly arrest in the first G1 phase immediately after cytokinesis failure, cancer cells progress through at least one cell cycle with doubled chromosome and centrosome numbers (Kuffer et al., 2013). Thus, USP28 loss improves proliferation of tetraploid cells by diminishing the negative impacts of tetraploidy in subsequent cell cycles.

To investigate how USP28 affects proliferation of tetraploids, we identified proteins interacting with USP28 specifically after cytokinesis failure, among them, most prominently, MDC1 and NuMA1 (Fig 5 A). NuMA1, a specific USP28 interactor after cytokinesis failure, plays multiple roles to ensure mitotic spindle integrity. It is recruited to minus-end of microtubules and acts as an adaptor for dynein and dynactin to allow spindle pole organization by generating pulling forces that control the spindle position (Okumura et al., 2018). Although loss of USP28 did not affect the protein levels of NuMA1 nor its localization, it increased the clustering of extra centrosomes and thus reduced multipolar mitoses. High levels of NuMA1 in tumor cells lead to high rates of multipolar mitoses and conversely, depletion of NuMA1 improves clustering of multiple centrosomes (Quintyne et al., 2005). As USP28 and NuMA1 likely interact in mitotic cells and USP28 depletion did not have an impact on NuMA1 protein levels, our data suggest a new ubiquitin-dependent mechanism that might be involved in control of clustering of extra centrosomes. Interestingly, recent evidence suggests a function of NuMA1 also in cellular response to DNA damage (Moreno et al., 2019). Future research will show whether this novel function of NuMA1 also contributes to cellular response to tetraploidy.

Next, we identified MDC1, a key mediator of the DNA damage response and replication checkpoint to interact with USP28 specifically in tetraploid cells. USP28 is a substrate for ATM in DNA damage response that stabilizes Chk2 and 53BP1 (Zhang et al., 2006). We observed increased phosphorylation of RPA32 and CHK1 that suggest replication stress in tetraploid cells and loss of USP28 diminished the signaling (Fig 6A,B). Thus, we considered the possibility that DNA damage response plays an important role in limiting proliferation of tetraploid cells. There is only scattered evidence suggesting that tetraploid cells suffer from increased DNA damage and the cause of DNA damage has not been clearly identified. One possibility is that the cells that do not arrested immediately after cytokinesis failure often undergo erroneous mitosis. Consequently, daughter cells inherit imbalanced chromosome number and become aneuploid, which may lead to replication stress and increased DNA damage (Passerini et al., 2016). Oxidative DNA damage also increases upon tetraploidization, which may also contribute to increased genomic instability (Kuffer et al., 2013). Finally, two recent publications showed that the two nuclei in binucleated cells after cytokinesis failure in human cell line or in Drosophila replicate asynchronously and accumulate DNA damage (Nano et al., 2019, Pedersen et al., 2016). Further investigation of how tetraploidy increases DNA damage will be important to understand how whole genome doubling contributes to evolution of cancer genomes.

## Material and Methods

### Cell lines and cell culturing conditions

HCT116 is a human colorectal carcinoma cell line, purchased from ATCC (No. CCL-247). HCT116-H2B-GFP were generated by transfection of WT cells with pBOS-H2BGFP construct (BD Pharmingen). HCT116 FUCCI were generated by transfection with a plasmid encoding the N-terminus of Cdt1 fused to mKO2 (an orange fluorescent protein) and is therefore present only in G1 phase (G1 sensor) and degraded in S, G2 and M phase by the SCF complex. The G2 sensor consists of the N-terminus of Geminin fused to mAG and is therefore degraded between anaphase and S phase by the APC/C complex (Sakaue-Sawano et al., 2008). RPE1-hTERT is a human retinal pigment epithelium cell line immortalized by telomerase expression (referred to as RPE1). RPE1 and RPE1-H2B-GFP cell lines were a gift from Prof. Erich Nigg (MPI Biochemistry, Martinsried) and Dr. Stephen Taylor (The University of Manchester, UK, Manchester Cancer Research Centre), respectively. DLD1 were provided by FB, CaCO2 were a kind gift from Prof. Axel Ullrich, MPI Biochemistry, Martinsried), HEK293 cell line was a kind gift from Prof. Stefan Kins (TU Kaiserslautern). Cells were cultured in Dulbecco’s Modified Eagle Medium GlutaMAX (DMEM) with addition of 10 % Fetal Bovine Serum (FBS) and 1 % Pen-Strep (50 IU/ml penicillin and 50 μg/ml streptomycin, PAA) in humidified cell culture incubator at 37°C with concentration of CO_2_ at 5 %. To passage the cells, cell culture dish was washed 2x with PBS, subsequently 0.25 % Trypsin-EDTA was added and incubated for 3 min in cell culture incubator. Cell culture medium was then added to inhibit trypsin, cells were collected, spun down and re-seeded.

### RNAi based screen for factors affecting the proliferaton of tetraploid cell

The HCT116 FUCCI (Fluorescent Ubiquitination-based Cell Cycle Indicator) cells were reverse-transfected with the esiRNA library in 384 well black glass bottom plates on Day 1. The next day DCD was added to a final concentration of 0.75 μM and 18 h later the DCD was washed out. 24 h later the cells were fixed with paraformaldehyde and staining with DAPI. Four fields *per* well were acquired by microscopy (Olympus ScanR High-Content Screening Station). The fluorescence intensity of the DAPI and the FUCCI signals of each nucleus was measured to establish the cell cycle profiles of cells expressing the FUCCI-G1, FUCCI-G2 or both FUCCI cell cycle probes, thus dividing the cells into six distinguished cell cycle classes 2CG1, 2CS, 4CG2, 4CG1, 4CS and 8CG2 according to their DNA content and the FUCCI sensor expression. Cells lacking FUCCI signal were excluded from the analysis. To account for systematic errors caused by batch effects and effects nested underneath due to the automatic liquid handling (8-channel dispenser and 96-channel washer) as well as plate edge effect, the absolute count of each of the six classes was corrected employing a linear model. The relative abundance was calculated and transformed into a Z*-score value for each of the six classes. The Z*-score transformation was performed for each cell cycle class by dividing the difference between its relative abundance in a particular well of a plate and the median of the whole plate by the median absolute deviation (MAD) (Zhang, 2006). Control wells transfected with esiRNA targeting either TP53 or KIFC1 as positive controls were excluded from the calculation of the median and MAD of the plate. Quantitative measures of four populations 2CG1, 4CG2, 4CG1 and 8CG2 were used to calculate a “Z-index” as a sum of the Z* scores of % 4CG2 and % 8CG2, from which the Z* scores of % 2CG1 and % 4CG1 were subtracted to reflect the proliferation of tetraploid cells. The primary screen was conducted in two technical replicates and the duplicate information was used to reduce the false negative discovery rate. The Z-index cutoff for candidates to score as a primary hit was set to 5.875. Using this strategy, we identified 432 genes that specifically increase the proliferation of tetraploids (TP53-like hits) among of the 16231 genes tested in the primary screen. A subset of 374 genes from the TP53-like category was selected for confirmatory screen, including the identified primary TP53-like hits, and excluding genes that either were identified as hits in previous cell cycle screens (Kittler et al., 2007; Neumann et al., 2010), or are located on the Y chromosome, which is not present in HCT116 cells, or were found previously not to be expressed in HCT116. The confirmatory screen was performed in black 96-well glass bottom plates in four technical replicates. Every assay plate contained four wells of renilla luciferase (R-LUC) and four wells targeting *TP53*, as negative and positive controls, respectively. The R-Luc wells were used for the Z*-score transformations. The TP53 wells (positive controls) separated well from the R-LUC wells; 56 out of 60 TP53 wells had a Z-index above 5.875 and the Z-index of the R-LUC controls was between −5.875 and 5.875 for 71 out of 72 wells. To confirm the primary TP53-like hits, every rescreened gene was tested against the R-LUC controls using the Dunnett’s multiple comparison test; we considered a primary hit as confirmed if the p-value was less than 0.1. Using this approach, 157 genes out of 374 primary hits were confirmed as TP53-like hits. The total confirmation rate was 42 %. For more details, see Supplementary notes.

### Generation of HCT116 H2B-GFP γ-tubulin-mRuby cell line

Human embryonic kidney cells (HEK293) were co-transfected with packing plasmid and γ-tubulin-mRuby plasmid using lipofectamine 2000 in Optimem. The medium was replaced with culture medium 24 hours post transfection. 24 h and 48 h later medium containing viral particles was collected and filtered. Next, HCT116 H2B-GFP cells were seeded in 6-well plate and infected using 100 μl and 600 μl supernatant per well with addition of Polybrene. After 48 hours, the cells were collected and seeded into 15 cm cell culture dish and kept in selection medium containing Zeocin 600 μg/ml. Individual clones were screened by fluorescence microscopy and those with γ-tubulin-mRuby expression were further cultured.

### Formation of tetraploid cells

Cells were treated with 0.75 μM actin depolymerizing drug dihydrocytochalasin D (DCD, Sigma) for 18 hours. Subsequently, the drug was washed out 3x using prewarmed PBS. Cells were further cultured in a drug-free medium for indicated time or immediately harvested for further experiments.

### Generation of CRISPR Cas9 knockout cell lines

First, Cas9 containing supernatant was produced using the same protocol as for γ-tubulin-mRuby. Next, HCT116 cells were transduced with Cas9 and, 48 hours later, the cells were reverse transfected with guideRNA and trackRNA in 6-well plates. The transfected cells were then seeded at 100 cells in 15 cm dish to obtain single clones that were picked and tested for protein expression by immunoblotting. The guide RNA were purchased from Dharmacon: USP28 (CR-006076-01-0002, CR-006076-02-0002, CR-006076-03-0002), SPINT2 (CR-010210-01-0002, CR-010210-03-0002, CR-010210-04-0002) and tracking RNA (U-002000-50). The arising mutations were validated by sequencing the specific clones.

### Transfection

For reverse siRNA transfection, the transfection mix (50nM siRNA mixed with lipofectamine 2000 and Optimem medium) was prepared and pipetted into 6-well plates. Cells were then seeded onto the plate in DMEM without antibiotics and incubated for 24 hours. Subsequently, the transfection medium was replaced with standard cell culture medium.

For plasmid transfections, the cells were seeded in 6-well plate and transfected using a forward protocol. Transfection mix consisting of titrated plasmid, lipofectamine 2000 and Optimem was added to the cells kept in DMEM without antibiotics. After 24 hours of incubation, transfection medium was replaced with cell culture medium.

### Proliferation assay

24 hours after siRNA transfection, the cells were treated with 0.75 μM DCD for 18 hours. After the drug washout, the cells were cultured in standard conditions for indicated time (0h, 4h, 8h, 24h, 30h, 48h). The cells were then pulse-labelled with 5-ethynyl-2’deoxyuridine (EdU) for 30 min, harvested and subjected to flow cytometric analysis.

### HDAC inhibition

The cells were seeded in 6-well plates and treated with 0.3 μM TSA for 6 hours and subsequently with 1 μM DOX for 16 hours, with DOX only, or left untreated. Harvested cells were pelleted and processed for immunoblotting.

### Mitogenic signaling inhibition and activation

To asses mitogenic signaling activation, the cells were transfected with siRNA and left to recover for 48 hours, then serum-starved overnight. Medium with serum was then added and cells were harvested in indicated timepoints and processed for western blotting.

### RT-qPCR

To assess the mRNA levels after knock downs, the cells were treated with siRNA for 24 hours, then cultured in standard conditions for 3 days or treated with 0.75 μM DCD and cultured 24 hours after the drug washout and subsequently collected. Total mRNA was isolated using a Qiagen mRNeasy mini kit according to manufacturer’s protocol. Next, reverse transcription using Anchored–oligo (Vigano et al.) and Roche Transcriptor First Strand cDNA synthesis Kit (Cat no. 04 379 012 001) was performed to obtain cDNA. Quantitative PCR was performed using specific primers and SsoAdvanced Universal SYBR Green Supermix (Bio-Rad, USA). Melting curve analysis was performed to confirm the specificity of amplified product. Each sample was spiked with TATAA Universal RNA Spike II control (TATAA Biocenter AB, Sweden). mRNA expression of each sample was normalized to control housekeeping gene *RPL30*.

### Fixed-cells imaging

Cells were seeded and treated when required in a glass-bottom 96-black well plate. The cells were then fixed using ice cold methanol, permeabilized with 3 % Triton X 100 in PBS and blocked in blocking solution (5 % Fetal Bovine Serum + 0.5% Triton X 100 + 1% Na_3_N in PBS). Subsequently, the plate was incubated with primary antibody overnight in 4°C.

Cells were washed 3x with PBS and incubated with secondary antibody and DAPI or Vybrant DyeCycle™ Green for 1 hour in RT. Before imaging, cells were washed 4x with PBS.

Imaging of fixed cells was carried out on a spinning disc system comprising of inverted Zeiss Observer.Z1 microscope, Plan Apochromat 63x magnification oil objective, 40x magnification air objective or 20x magnification air objective, epifluorescence X-Cite 120 Series lamp and lasers: 473, 561 and 660 nm (LaserStack, Intelligent Imaging Innovations, Inc., Göttingen, Germany), spinning disc head (Yokogawa, Herrsching, Germany), CoolSNAP-HQ2 and CoolSNAP-EZ CCD cameras (Photometrics, Intelligent Imaging Innovations, Inc., Göttingen, Germany). SlideBook software (Intelligent Imaging Innovations, Inc., Göttingen, Germany) was used for image acquisition and analysis.

### Live-cell imaging

Cells expressing H2B-GFP were seeded in a 96-well plate at 30 000 cells per well in standard cell culture medium. Subsequently, the cells were treated with 0.75 μM DCD to induce tetraploidization. After the treatment, the cells were washed with prewarmed PBS and FluoroBrite medium was added to proceed with live-cell imaging. Imaging was performed using an inverted Zeiss Observer Z1 microscope (Visitron Systems) equipped with a humidified chamber (EMBLEM) at 37°C, 40% humidity, and 5% CO_2_ using CoolSNAP HQ2 camera (Photometrics) and X-Cite 120 Series lamp (EXFO) and Plan Neofluar 20x, or 10x magnification air objective NA 1.0 (Zeiss, Jena, Germany). Cells were imaged for 48 hours with 8-min time-lapse. Images were analyzed using Slidebook (Intelligent Imaging Innovations, Inc., Goettingen, Germany) and ImageJ (National Institutes of Health).

### Cell lysis and protein concentration measurement

Pelleted cells were lysed in RIPA buffer with protease inhibitor cocktail (Pefabloc SC, Roth, Karlsruhe, Germany), then sonicated by ultrasound in a water bath for 15 min. Cell lysate was spun down at 13600 rpm for 10 min at 4°C on a table-top microcentrifuge (Eppendorf, Hamburg, Germany). 1 μl was used to determine protein concentration using Bradford dye at 595 nm wavelength. Subsequently, the lysates were mixed with 4x Lämmli buffer with 2.5% ß-mercaptoethanol and boiled at 95°C for 5 min.

For fractionation, the cells were lysed in 0.1% ice cold NP-40, then spun down. Supernatant was transferred to a separate tube as the cytoplasmic fraction and the pellet was processed using RIPA buffer to obtain the nucleoplasmic fraction. Lysates were further processed as described.

### SDS-PAGE and immunoblotting

Prepped cell lysates were separated by SDS-PAGE using 10% or 12.5% gels. Protein size was estimated using the PrecisionPlus All Blue protein marker (Bio-Rad, USA). Gels were incubated in Bjerrum Schafer-Nielsen transfer buffer and proteins were transferred to a water-activated nitrocellulose membranes (Amersham Protran Premium 0.45 NC, GE Healthcare Life Sciences, Sunnyvale, USA) using semi-dry transfer (Trans-Blot® Turbo™, Bio-Rad, USA). Membranes were stained in Ponceau solution for 5 min and scanned to be used as a loading control. Next, membranes were blocked in 5% −10% skim milk in TBS-T (Fluka, Taufkirchen, Germany) for 1 h in RT. After blocking, membranes were incubated in respective primary antibodies diluted in 1 % Bovine Serum Albumin (BSA) or 5 % skim milk overnight at 4°C with gentle agitation. Further, the membranes were rinsed 3 x 5 minutes with TBS-T, incubated 1 h in RT with HRP-conjugated secondary antibodies (R&D Systems), and followed by rinsing 3x 5 minutes with TBS-T. Chemiluminescence was detected using ECLplus kit (GE Healthcare, Amersham™) and Azure c500 system (Azure Biosystems, Dublin, USA). Protein band quantification was carried out using ImageJ (National Institutes of Health, http://rsb.info.nih.gov/ij/). All used antibodies are listed in Supplementary table 5.

### Chromatin immunoprecipitation (CHIP)

Cells were treated with siRNA and 0.75 μM DCD (18 h treatment and release for 24 h or 48 h) or 1 μM DOX (16h) was added one day after transfection. Treatments were arranged in a way that all samples were collected simultaneously. The experiment was performed using SimpleChIP^®^ Enzymatic Chromatin IP (Magnetic Beads) kit (Cell Signaling, #9003) according to manufacturer’s protocol. Briefly, the cells were treated in cell culture dish with 37% PFA to fix and crosslink the proteins to the DNA. Cells were then collected, incubated with micrococcal nuclease and sonicated. Digested chromatin was subsequently incubated with an antibody (anti-p53, anti-H3, Normal Rabbit IgG) and incubated overnight at 4°C with rotation. Next, magnetic beads were added to each immunoprecipitation sample and incubated for 2 hours, followed by washing steps and elution. DNA was purified and used for RT-qPCR.

### Co-immunoprecipitation

Untreated and DCD-treated cells were collected and lysed in lysis buffer (25mM Tris-HCl, pH 7.4, 150mM NaCl, 1mM EDTA, 1% NP-40, 5% glycerol). Magnetic beads (SureBeadsTM Magnetic Beads, Bio-Rad, USA) were conjugated with antibody (anti-USP28, anti-SPINT2, anti-Flag) according to manufacturer’s protocol. Next, cell lysates were incubated with the beads overnight, 4°C, washed 2 times with washing buffer I (25mM Tris-HCl, pH 7.8, 500mM NaCl, 0,5% Triton-X 100) and 3 times with washing buffer II (10mM Tris-HCl, pH 7.8, 150mM NaCl) and eluted with 2x Lämmli. Samples were loaded onto the SDS page gel and either processed further to confirm pull-downs by immunoblotting or to detect interactors by mass spectrometry.

### FACS analysis of proliferation

Harvested cells were spun down and fixed using the Fix-Perm buffer. Afterwards, the samples were incubated with Click-it reaction mix for 30 min in the dark followed by 30 min incubation with anti-cyclin B antibody. Next, the samples were incubated for 30 min with secondary antibody. Between each incubation, the cells were washed 3 times with Perm-Wash buffer and spun down. After the incubation with secondary antibody, the cells were resuspended in PBS RNase (RNase Zap, Invitrogen, Carlsbad, USA) solution with 4’,6-Diamidino-2-Phenylindole, Dihydrochloride (DAPI) and measured using Attune Nxt acoustic focusing cytometer (Life Technologies, Carlsbad, USA).

### Mass spectrometry identification of interacting proteins

Mass spectrometry identification of USP28 was performed by Nagarunja Nagaraj at the Mass Spectrometry Core Facility at the MPI Biochemistry, Martinsried, Germany, as previously described (Tyanova, 2016). The identified proteins generated by MaxQuant V were uploaded to Perseus V 1.6.2.3 (Tyanova, 2016). Site only, reverse, and contaminant peptides were removed from the dataset and missing values were imputed using a normal distribution. Invalid values were then excluded. The volcano plot function was used to identify proteins that were significantly altered using a t-test with a false discovery rate of 0.05 and an S_0_ of 0.1.

### SPINT2 and CDKN1A tissue sample mRNA expression correlation

Log-transformed gene expression values of SPINT2 and p21 gene CDKN1A quantified using RNA-Seq by Expectation Maximization (RSEM) (Li, 2011) with data from The Cancer Genome Atlas (TCGA) (http://cancergenome.nih.gov/). The samples comprise 285 primary colon adenocarcinoma and 41 normal tissue samples taken from colon cancer patients. Pearson’s correlation coefficient rho has been used for gene expression correlation testing.

## Acknowledgement

We thank Isabell Kirchner and Robin Roth for excellent technical support. We thank Stephen Elledge (Harvard Medical School, Boston, MA) for providing the USP28 carrying vectors, Lotte Vogel (University of Coppenhagen, DK) for providing the SPINT2 carrying vector and Holger Bastians (University of Gottingen, Germany) for providing the γ-tubulin-RFP carrying vector. This work was supported by the European Union through an FP7 Marie Curie Innovative Training Network grant, PloidyNet (to ZS) and by DFG FOR2800 (to ZS and MK). The results presented in Fig 3H are based upon data generated by the TCGA Research Network: https://www.cancer.gov/tcga.

## Author contributions

The genome wide screen and its evaluation was performed by C.K., D.K., M.T. and F.B. K.S.-T. and S.V.B performed the follow-up experiments. K.K. performed the proteomics analysis of the pull down experiments, J.-E. B. and M.K. conducted the meta-analysis of cancer genomics data. Z.S. conceived the study, C.K., K.S.-T., S.V.B. and Z.S. designed and analyzed the experiments, Z.S. wrote the manuscript, all authors discussed and edited the manuscript.

## Tables and their legends

Supplementary table 1 Summary results of the screen

Supplementary table 2 Pathway analysis of the identified candidates

Supplementary table 3 Comparison with other screens’ results

Supplementary table 4 Results of the mass spectrometry of USP28 pull down

Supplementary table 5 List of used antibodies

## Expanded View Figure legends

**Expanded View 1** Validation of selected candidates

(A) Cell cycle profile after a release from DCD in HCT116.

(B) Efficiency of siRNA depletion evaluated by q-RT-PCR.

(C) Proliferation evaluated as fold changes of EdU incorporation in diploid cells after depletion of respective factors. siNT - non targeting control.

(D) Example of an immunoblot for validation of the SPINT2 knock out. Sequencing of the SPINT2 confirmed a short deletion within the exon 2. The knock out cell lines also lost the ability to induce p21 expression. Four clones were isolated: S15, S25, S28 and S40.

(E) Example of an immunoblot for validation of the USP28 knock out. Three clones were isolated: U22, U35 and U46.

(F) Examples of cell cycle profiles in the knock out cell lines 24, 30 and 48 after induced cytokinesis failure.

(G) Validation of the overexpression of USP28 and SPINT2, respectively, in knock out cell lines. Empty – an empty control vector, WT: a vector carrying the wt gene; C171A - vector carrying the mutant version lacking the deubiquitinase activity. DRB - doxorubicin.

**Expanded View 2** Activation of PIDDosome and Hippo pathway in tetraploid HCT116 cells

(A) MDM2 and CAS2 cleavage in HCT116 upon treatments with DCD, centrinone and reversine.

(B) MDM2 and CAS2 cleavage in HCT116 upon depletion of p53, USP28 and SPINT2.

(C) Phosphorylation of LATS2 in HCT116 cells untreated or treated with DCD for 18 h.

(D, E) Immunoblot and quantification of YAP nuclear enrichment in HCT116 upon DCD treatment. Mean and SEM of three independent experiments is shown.

**Expanded View 3** Activation of the cMET pathway

(A) Schematic depiction of the cMET pathway.

(B) Representative immunoblot of cMET downstream kinases upon realease from starvation.

(C) Quantification of AKT phosphorylation and CDKN1A expression upon realease from starvation. Means of three experiments + SEM are shown.

(D, E) Immunoblot and quantification of cMET target c-Myc upon release from DCD. Mean and SEM of three independent experiments are shown, paired T test was used for statistical evaluation; * p<0.05.

**Expanded View 4** USP28 and NUMA1 in response to tetraploidy

(A) Fraction of cells in S phase upon depletion of USP28, p53 and 53BP1. Means of three independent experiments + SEM are shown, normalized to untreated control (ctrl). T test was used for statistical evaluation; * p< 0.05; ** p < 0.01; *** p< 0.001.

(B) Time in mitosis in tetraploid HCT116 and in knock out cell line lacking USP28 (U22). T test was usedfor statistical evaluation; * p< 0.05; ** p < 0.01; *** p< 0.001.

(C) Localization of NUMA1 and USP28 during mitosis.

(D) Representative immunoblot of NUMA1 and USP28 upon respective depletion.

(E) Quantification of NUMA1 and USP28 levels upon siRNA mediated depletion. Means of three experiments + SEM are shown. T test, * p<0.05.

**Expanded View 5** DNA damage and checkpoint response in tetraploid cells.

(A) Quantification of CHK1 levels after DCD induced tetraploidization in HCT116 and in cells lacking USP28 (U35). upon siRNA mediated depletion. Means of three experiments + SEM are shown. T test - no significant difference. (B) RPA32, as in (A).

